# Noninvasive sampling for comparisons of wildlife microbiomes may be more reliable than sampling trapped animals

**DOI:** 10.1101/2022.02.17.480865

**Authors:** Sondra Turjeman, Sasha Pekarsky, Ammon Corl, Pauline L. Kamath, Wayne M. Getz, Rauri C. K. Bowie, Yuri Markin, Ran Nathan

## Abstract

In ecological and conservation studies, responsible researchers strive to obtain rich data while minimizing disturbance to wildlife and ecosystems. We assessed if samples collected noninvasively can be used for microbiome research, comparing microbiota of noninvasively collected fecal samples to those collected from trapped common cranes at the same sites over the same periods. We found significant differences in microbial composition (alpha and beta diversity), which were not accounted for by noninvasive samples’ exposure to soil contaminants, as manually assessed by comparing differentially abundant taxa. They could result from trapped birds’ exposure to sedatives. We conclude that if all samples are collected in the same manner, comparative analyses are valid, and noninvasive sampling may better represent host microbiota because there are no trapping effects. Experiments with fresh and delayed sample collection can elucidate effects of environmental exposures on microbiota. Further, stressing or sedation may unravel how trapping affects wildlife microbiota.

## Introduction

Microbiome research, in non-model species, has proliferated in the past decade. A number of studies on diverse species have shown that endogenous microbiota can be used to assess host state (Peixoto et al., 2021). Microbiota manipulation (fecal microbiota transplant), already a successful tool in treating some human maladies, is being examined in wildlife as a treatment option (Guo et al., 2020; Niederwerder, 2018). Studies have suggested the presence of a core microbiota that has co-evolved with the host species (Risely, 2020) and have also revealed that much of a host’s microbiota is relatively dynamic in composition, with shifts in response to environment, diet, or other changes in physiology and life-history. Thus, there are many potential applications of microbiota profiling in conservation ecology (Trevelline et al., 2019). Populations can be monitored for overall health (Peixoto et al., 2021) and for the presence/absence of specific pathogens (Choi et al., 2021). Compositional comparative analyses—or even interventions—of captive and wild populations microbiota (Gibson et al., 2019; Oliveira et al., 2020) can be made prior to reintroductions. Recently, even reproductive health and potential breeding success of wild animals was assessed from microbiota profiling (Comizzoli & Power, 2019).

While microbiota profiling can be a powerful tool in humans and animals alike (Turjeman & Koren, 2021), trapping and handling animals poses challenges to conversation research. Often, animals are difficult to trap, efforts can be cost- and resource-prohibitive, and many species do not withstand the stress of trapping well (Beja-Pereira et al., 2009). Further, stress associated with trapping and handling can affect the microbiome (Collins & Bercik, 2009) even if it does not seem to visibly harm the target species, thus skewing findings. In other realms of conservation biology—population genetics, evaluation of mating systems, species surveys and counts—noninvasive sampling and environmental sampling combined with molecular tools have provided powerful means to study animals while keeping disturbances to a minimum (e.g. (Russello et al., 2015). Here we assess the utility of non-invasive sampling in microbiome studies by comparing the composition of fecal samples freshly defecated from trapped common cranes (*Grus grus*) to noninvasively collected samples of different cranes collected in the same area. While we detected non-negligible differences in the microbial composition of feces collected using the two sampling methods, we also found that the total number of represented taxa for each method were both high and similar, suggesting that if all samples are collected in the same noninvasive manner, comparative analyses between populations or across time are likely valid. Of note, our noninvasive method is truly noninvasive – handing of animals is not required for any parts of the method unlike unlike (Knutie & Gotanda, 2018) and (Pannoni et al., 2022).

## Methods

### Sample collection

We obtained samples for this project as part of a wider study of common crane movement, foraging, and microbiome (Pekarsky et al., 2021). During the pre-migration period (early fall), cranes stage in western Russia (Ryazan area; 54°56′N, 41°02E) for several weeks before the onset of migration (Johnsgard, 1983; Leito et al., 2015). We trapped 27 cranes during this period in 2017 using bait mixed with alpha-chlorolose, a routine oral sedation technique associated with low morbidity and mortality rates (Hartup et al., 2014). After trapping, birds were hooded, banded, GPS-tagged, and measured. Then hoods were removed, and birds were held under constant supervision until complete recovery from sedation. When they defecated onto the ground (usually soon before recovering from sedation and flying away), we sampled the inner portion of the fresh feces using sterile cotton swabs, stored it immediately in 95% EtOH at −20°C, and then transferred to −80°C for long-term storage. Average time elapsed between trapping and sample collection was 405 min (range 39-1278 min, based on when GPS data-loggers recorded locations beyond the trapping site). In parallel, we collected 37 fecal samples from the ground following observations of birds defecating in fields near the trapping site in the two weeks prior to and the two weeks during trapping. When a flock of cranes was observed, we scattered them by approaching the flock and then collected samples from individual droppings (>30cm apart) as above. We assume that samples were not from trapped birds, as GPS data suggested tagged birds were not present at the immediate site of sampling, and there were hundreds of birds in the staging area during this period, but the possibility cannot be ruled out completely. If we inadvertently sampled a subset of birds with both methods, we might observe slightly greater overlap in the two sampling-methods’ microbiota profiles, but this should not confound results. Post-sampling, samples were handled and stored identically.

### Microbial sample sequencing and pre-processing

Sample extraction and processing for 16S rRNA metabarcoding is described in Pekarsky et al. (2021). Briefly, we PCR-amplified the V4 region of the 16S rRNA in triplicate for sequencing on an Illumina MiSeq. Data was processed in R v.4.1.1 to identify amplicon sequence variants (ASVs) as described in Pekarsky et al. (2021). In total, 64 samples were sequenced for this study. We used the package *decontam* (Davis et al., 2018) to identify and remove 27 contaminant ASVs (prevalence method, threshold: 0.5) based on three negative controls and two PCR blank samples sequenced in the same run. We removed sequences that could not be assigned to a phylum and those classified as mitochondria. We compared read depth between sampling methods using, as appropriate, *t*, Mann-Whitney *U*, and Kolmogorov-Smirnov tests. The median number of reads per individual across all samples following filtering was 14,397.5 (range: 5,337-25,670).

### Comparison of Microbiota

Analyses, unless otherwise specified, were performed using the *phyloseq* (McMurdie & Holmes, 2013) and *microbiome* (Lahti & Shetty, 2019) R packages. After examining rarefaction curves, we rarefied data to 8,000 reads to optimize the trade-off between read depth and sample size (3 samples lost; final read depth 8,044; *phyloseq*, seed: 999). We compared alpha diversity of the microbiota in fecal samples from trapped birds to that of birds sampled noninvasively using Faith’s phylogenetic diversity index (*btools*; (Battaglia, 2021) and raw observed richness with Mann-Whitney *U* tests. Differences in community composition were measured using both weighted and unweighted UniFrac measures and compared with PERMANOVAs. Within-group dispersion was assessed with the *vegan* package (Oksanen et al., 2018). Differentially abundant taxa were identified with *ANCOMBC* (Lin & Peddada, 2020). We used a minimum prevalence filter of 10% and an FDR threshold of 0.05 when identifying significantly differentially abundant taxa. A heatmap based on significantly different genera with an absolute log-fold-change > 0.58 was produced using *pheatmap* (Kolde, 2019). After identifying core microbiota for samples from each of the sampling methods, using a minimum threshold of 10% prevalence, we used *eulerr* (Larsson & Gustafsson, 2018) to generate a Venn diagram.

## Results

Analyses were based on 61 samples following rarefaction, 25 from trapped birds and 36 noninvasive samples. There was no difference in mean sequencing depth or sequencing depth distribution between the two sampling methods (*t*-test: *p*=0.17; Kolmogorov Smirnov: *p*=0.13; Fig. 1). Following rarefaction to 8,044 reads, comparison of alpha diversity using Faith’s PD (Mann-Whitney: *p*=0.0009) and observed richness (Mann-Whitney: *p*=0.0003) revealed that fecal samples from trapped birds consistently had greater alpha diversity than those sampled noninvasively from free-foraging cranes (Fig. 2a, b). There were significant differences in sample composition (beta diversity) of crane microbiota between the two methods, assessed with both weighted (PERMANOVA: *p*<0.0001; Fig. 2c) and unweighted (PERMANOVA: *p*<0.0001; Fig. 2d) UniFrac. Notably, community dispersion using the weighted measure was significantly greater among trapped birds (betadisper: *p*=0.034; Fig. 2c), but this pattern was not preserved with the unweighted metric (betadisper: *p*=0.29; Fig. 2d).

**Figure 1.**
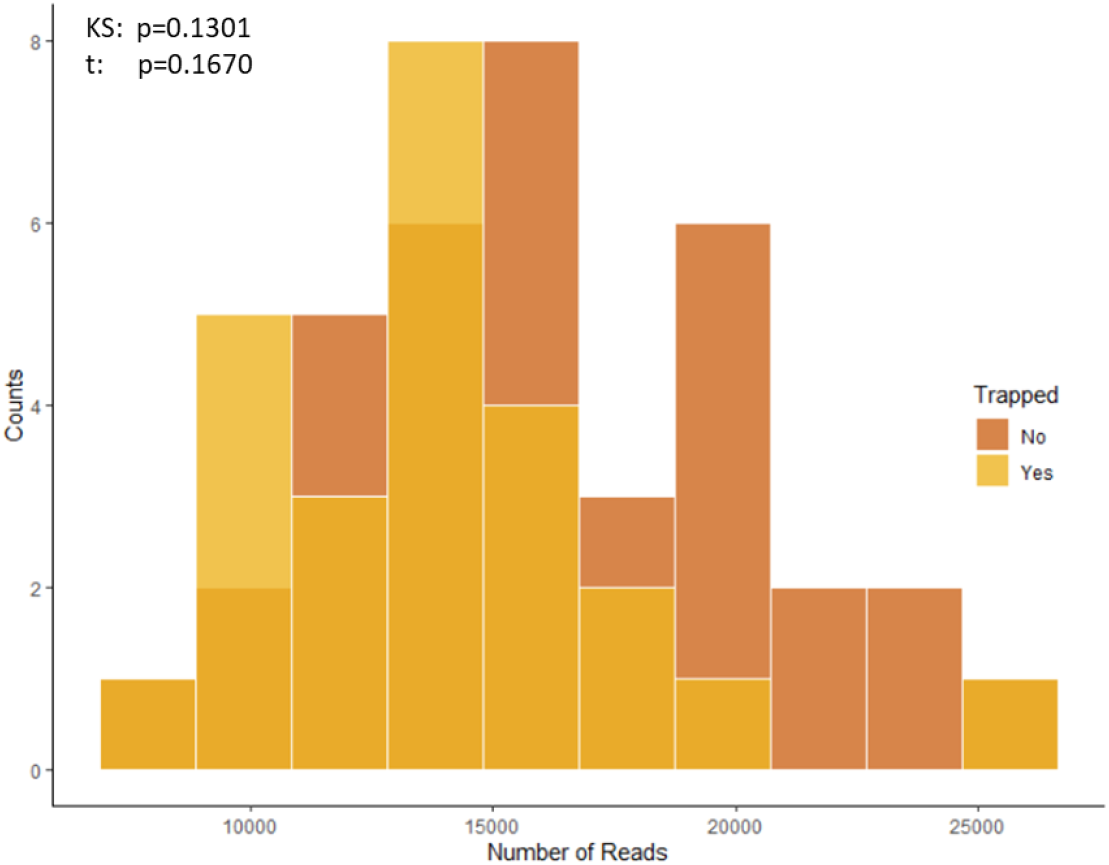
Sequencing depth for each sampling method. No differences in sequencing depth were found between samples collected from trapped birds versus those collected noninvasively from free-ranging birds.

**Figure 2.**
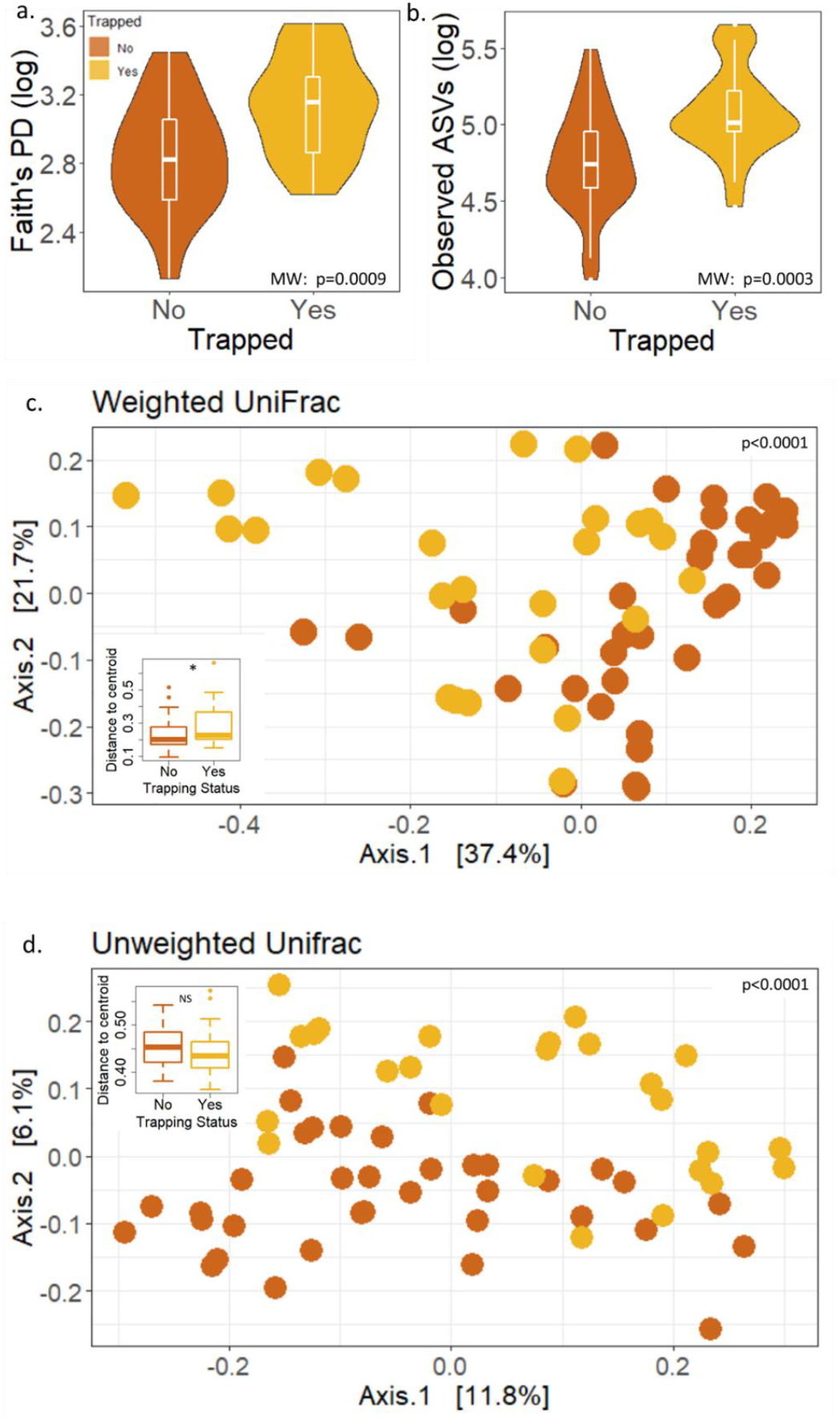
Comparison of microbiota for each sampling method. Samples collected from trapped birds had significantly higher diversity (a) and richness (b) compared to non-invasive samples. Furthermore, the two methods resulted in significantly separated communities in the PCoA space when using both weighted (c) and unweighted (d) UniFrac metrics. Dispersion of the groups (insets) was significantly different when using the weighted (c) but not unweighted (d) method, with trapped bird samples showing a more dispersed cluster of microbiota than noninvasive samples. Boxplots in violins: boxes represent medians with 1^st^ and 3^rd^ quartiles and whiskers represent maxima and minima.

Visual examination of community composition at the phylum level supports our findings of differences between the microbiota of trapped and noninvasively sampled cranes (Fig. 3a), and differential abundance analysis identified differentially abundant taxa, both at the phylum (Fig. 3b) and genus (Fig. 3c) leves. We examined the genus-level differences in light of both within-group relative abundance and within-group prevalence (Fig. 3c,d, Table S1) and found that only four of the differentially abundant taxa had a relative abundance greater than 0.1%. Similarly, less than one third of significant genera had a within-group prevalence above 90%. In total, we identified 28 different phyla in our samples, of which, 10 (36%) differed between the groups in their relative abundances. Of the 444 genera identified, relative abundances for only 23 (6%) differed. When comparing a “core” of genera found in at least 10% of samples per group (a combined 183 genera), we found that ca. 60% overlapped between sampling methods, whereas ca. 7% were only found in the noninvasive samples and ca. 32% were only found in trapped samples (Fig. 3e). We also examined the effect of various minimum relative abundance thresholds (weighted UniFrac: 0.1%: *p*<0.0001, 1%*p*<0.0001) and ran analyses at the genus level (weighted UniFrac: *p*<0.0001). In all cases, the different sampling methods resulted in significant differences in microbiota composition. Further, when examining the microbiota of the samples collected directly from trapped birds, we did not find evidence of age (weighted UniFrac: *p*=0.98) or sex (weighted UniFrac: *p*=0.44) stratification in the trapped birds’ microbiota.

**Figure 3.**
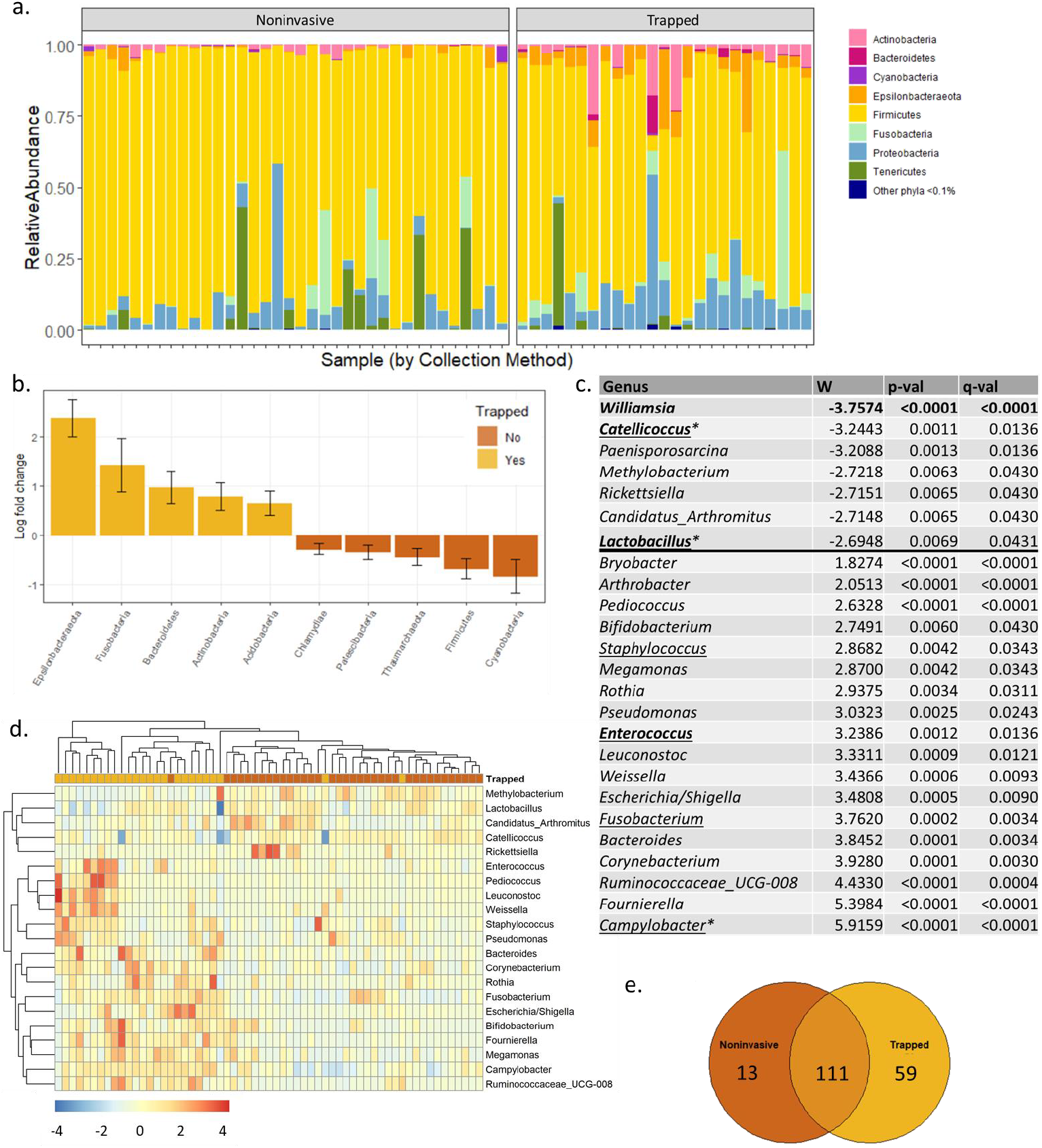
Differentially abundant taxa. We examined differentially abundant taxa between the microbiota of trapped and noninvasively sampled free-ranging cranes at the phylum (a,b) and genus (c) levels. Individual bar plots (a) suggest sampling method-specific differences at the phylum level, confirmed by ANCOM-BC (b). A positive log-fold-change denotes bacterial taxa relatively increased among trapped samples and a negative log-fold-change denotes those relatively increased in noninvasive samples. Only significant taxa, following FDR correction, are shown. At the genus level (c), we found 23 significant taxa. A positive W denotes the abundance of genera that increased in the trapped group, and a negative W denotes the abundance of genera that increased in the non-invasive group. Genera in bold are found in at least 90% of the noninvasive samples and those underlined are found in at least 90% of trapped bird samples. * denotes mean relative abundance of >0.1%. A heat map of the significant taxa (d) shows clustering of the noninvasive samples together based on differentially abundant genera. The color bar denotes normalized, standardized abundances (from ANCOM-BC). Only genera with a log-fold-change >0.58 are included. In total, we found (e) 111 shared genera when using a minimum threshold of 10% (for a given sampling method and 13 unique genera for the noninvasive samples vs. 59 unique genera for the trapped samples.

## Discussion

Despite differences between the microbiota from trapped and noninvasively sampled cranes, our results indicated that both methods can be used as a source of microbial matter for 16S rRNA metabarcoding. We did not find differences in sampling depth across sample types. We found consistent differences, though, in bacterial richness and composition, with the samples from trapped birds exhibiting significantly higher richness and dispersion in overall composition when considering both bacterial lineages and their abundances. We thus conclude that the two sampling methods can yield different microbiota profiles for the same population.

Overall, 16S rRNA sequencing of the two sample types resulted in similar sequencing depths and a comparable number of ASVs (trapped: 1403, noninvasive: 1272). Of the prevalent genera (a subset of ASVs shared by >10% of birds), 60% were shared across the sampling methods, suggesting a core microbiota can be successfully identified using either method. This is impressive given the high intra-individual diversity previously reported for common cranes (Pekarsky et al., 2021). Similarly, most of the differentially abundant genera were low-abundance taxa, again suggesting that either method can be used to identify many of the most common core microbial taxa. Removing rare taxa and rerunning analyses did not change our results, suggesting that while we can define a coarse core microbiota, the two sampling methods differ for some relatively common taxa (1% relative abundance, genus-level), which suggests that variation between the methods are not just due to random differences in sampling of rare taxa.

We next considered the differentially abundant taxa to better understand potential effects of sampling methods on microbial community composition. Of the 23 differentially abundant genera, only seven were found in >90% of trapped bird samples and four were found in >90% of noninvasive samples; thus, the differentiating taxa are neither highly prevalent nor extremely conserved. Looking at the overall abundances of these taxa, the picture is similar: only three genera have relative abundances >0.1%. Thus, most of the differentiating taxa appear to be somewhat rare. Rare taxa, though, can have important roles in the overall microbiota both in their specific functionalities and in maintaining overall gut and microbial homeostasis (Banerjee et al., 2018). We therefore conclude that our differential abundance analysis, together with the consistent differences in alpha- and beta-diversity, demonstrates that different sampling methodology may capture different microbiota. While we may be able to define common and even temporally stable (in cases of longitudinal sampling) core microbiota, it could be difficult to reach conclusions regarding the ecological, functional, and host-adapted cores (Risely, 2020). Our findings contrast with those of a recent study examining trapped and caged small mammal microbiota, which concluded that despite some differences in specific taxa, sampled communities were relatively similar. There were no differences in comparisons of alpha diversity or unconstrained comparisons of beta diversity between mice sampled after trapping and those sampled in their home cages (Čížková et al., 2021).

To determine if differences in the noninvasive samples’ microbiota were derived from extended exposure to aerobic conditions or contaminants, we manually examined functionality of the differentially abundant genera (Fig. 3C) over-expressed in noninvasive samples. *Catellicoccus* is a facultative anaerobe that has previously been found in gull feces (Koskey et al., 2014). Similarly, *Methylobacterium* and *Candidatus arthromitus* have also been found in birds (Escallón et al., 2019) and other animals (Snel et al., 1995), though the former is also found in soil and water and could be a contaminant. *Paenisporosarcina* is an aerobic genus that previously isolated from soil and *Williamsia* is also an opportunistic aerobe. Interestingly, *Lactobacillus*, an anaerobic genus, was enriched in the noninvasive samples, though there were more lactic acid bacteria and anaerobes among the significantly differentially enriched genera of the trapped samples. In contrast, we did find two genera overrepresented in the trapped samples, *Bryobacter* and *Arthrobacter*, that are typically found in soils. Together, these findings suggest that while the method of collecting a fecal sample from the middle of the dropping may help to reduce environmental contaminants and proliferation of (opportunistic) aerobic bacteria, it is not perfect. The last enriched genus among noninvasive samples, *Rickettsiella*, is associated with tick-borne diseases and ectoparasites sometimes found on migratory birds (Cerutti et al., 2018), and it is unclear why this specific taxon proliferated. We did not identify arthropods in noninvasive samples either during collection or sample processing in the lab.

Because soil contamination did not appear to be a main driver of microbiota differences, we considered a second source of microbiota differentiation: stress associated with trapping and handling. In our case, specifically, birds were not only exposed to stress, but also to a sedative. Stress effects on microbiota have been examined in a number of animal species (Noguera et al., 2018; Stothart et al., 2016), and anesthesia in mice was also found to have rapid (4 hr) effects on microbiota composition (Serbanescu et al., 2019). Similarly, sedated birds may defecate less frequently, and time in the digestive tract is known to affect microbiota composition (Vandeputte et al., 2016). Of note though, none of the taxa overrepresented in our trapped crane samples were enriched in yellow legged gulls (*Larus michahellis*) experimentally implanted with corticosterone. Rather, several, *Pseudonomas* and *Campylobacter*, enriched in our trapped samples, were underrepresented in the corticosterone-implanted birds (Noguera et al., 2018). Similarly, comparing enriched taxa in our trapped bird samples to those overrepresented in high- and low-stress lines of Japanese quails (*Coturnix japonica*) did not reveal a clear pattern of association to either group (Lyte et al., 2021). Finally, *Ruminococcaceae*, enriched in our trapped samples, is typically associated with reduced stress, so their abundance and those of other low- or no-stress taxa could be an immediate effect of alpha-chlorolose used in trapping. Alpha-chloralose makes birds less aware of their surroundings and generally less alert or even puts them to sleep for 1 to 24 hours, after which they awaken and freely leave the capture area (Hayes et al., 2003). This capture method, suggested as effective in alleviating stress and lowering stress-related morbidity (Hartup et al., 2014), is applied through ingestion and thus might affect gut microbial composition. Birds defecated only shortly prior to recovery from sedation; thus, large differences in recovery time (39-1278 min) and possible difference in stress and residual sedative levels, may explain the larger differences in microbiota composition in the trapped group (weighted UniFrac dispersion) compared to the non-invasively sampled birds. We did not however find a correlation between alpha diversity (Faith PD) and time until departure from the trapping site (p > 0.05).

In summary, we found significant differences among microbial communities revealed by sampling trapped versus non-trapped common cranes in the same sites over the same temporal periods. Some differences are likely associated with soil contamination, but others seem to reflect effects of trapping and oral sedation. While sedation might be a taxa-specific practice in wildlife research, trapping has been broadly applied, and potential effects of trapping (with and without sedation) on microbiota composition should be considered. Our results also suggest that the potential contamination of noninvasive samples might not be substantial, supporting expanded use in conservation ecology. Further, while some animals are relatively easy to trap, other species require complicated, expensive, or time-consuming methods that might impede the ability to reach sufficient sample sizes (Sutherland et al., 2004), and trapping of wild animals may induce stress and endanger fitness, which is especially problematic for rare or endangered species (Blomberg et al., 2018; Dennis & Shah, 2012; Spotswood et al., 2012). Consequently, removing stress and other effects of interventions associated with trapping could not only promote animal welfare but also allow for sampling of a microbiome that presumably more closely represents that of the host under typical conditions.

Fecal microbiota characterization from noninvasively collected samples may not be perfect, but neither is that from trapped birds. Further, if all noninvasive samples are collected in the same manner, we believe that comparative microbiota analyses and monitoring will still be valid. The total number of represented bacterial taxa detected by each sampling method are high, suggesting that both methods provide informative microbiota data. When comparing noninvasive data from one population with that of another population, we anticipate that observed differences will be biologically relevant (Pekarsky et al., 2021), though environmental sampling should be considered to identify and remove site-specific contaminants.

Further investigations into how to best characterize host microbiota noninvasively would be useful because noninvasive sampling greatly improves research reach across study systems and reduces trapping-induced microbial changes. Controlled experiments with captive animals—collecting feces via swab, fresh following excretion, and at various increments following defecation—should shed light on how samples change post-defecation. This information could be used to correct microbiota characterizations made from noninvasively collected samples, allowing researchers to filter taxa that proliferated during the time to collection. It could be combined with findings from controlled stressing or sedating experiments in wild or semi-wild setting towards better understanding how stress effects the microbiota of non-model organisms. Regardless, we demonstrate the utility of noninvasive sampling of fecal samples as means of characterizing the microbiota of a large and difficult-to-trap bird species. We anticipate widespread use in comparative analyses, health surveys, and pathogen tracking of wild species. Most importantly, though, is sampling method consistency throughout a projects’ lifetime.

## Acknowledgements and Data

We acknowledge K. Postelnykh and K. Kondrakova help in fieldwork and trapping. We thank M.Lee-Folz, H. Newcombe, and L. Smith for assistance with laboratory work. In addition, we thank R. Efrat for highly pertinent feedback on early drafts of the manuscript and E. Sharon for help generating the heatmap from our results. All field procedures carried out in this study were approved by the Department of Environment of the Ryazan District, Russia (permit CK19-7154). The sequence data generated as part of this study will be deposited in the Sequence Read Archive upon acceptance of the manuscript. Final codes will be included in the final supplementary file after revisions.

---

**Supplementary Table S1.**
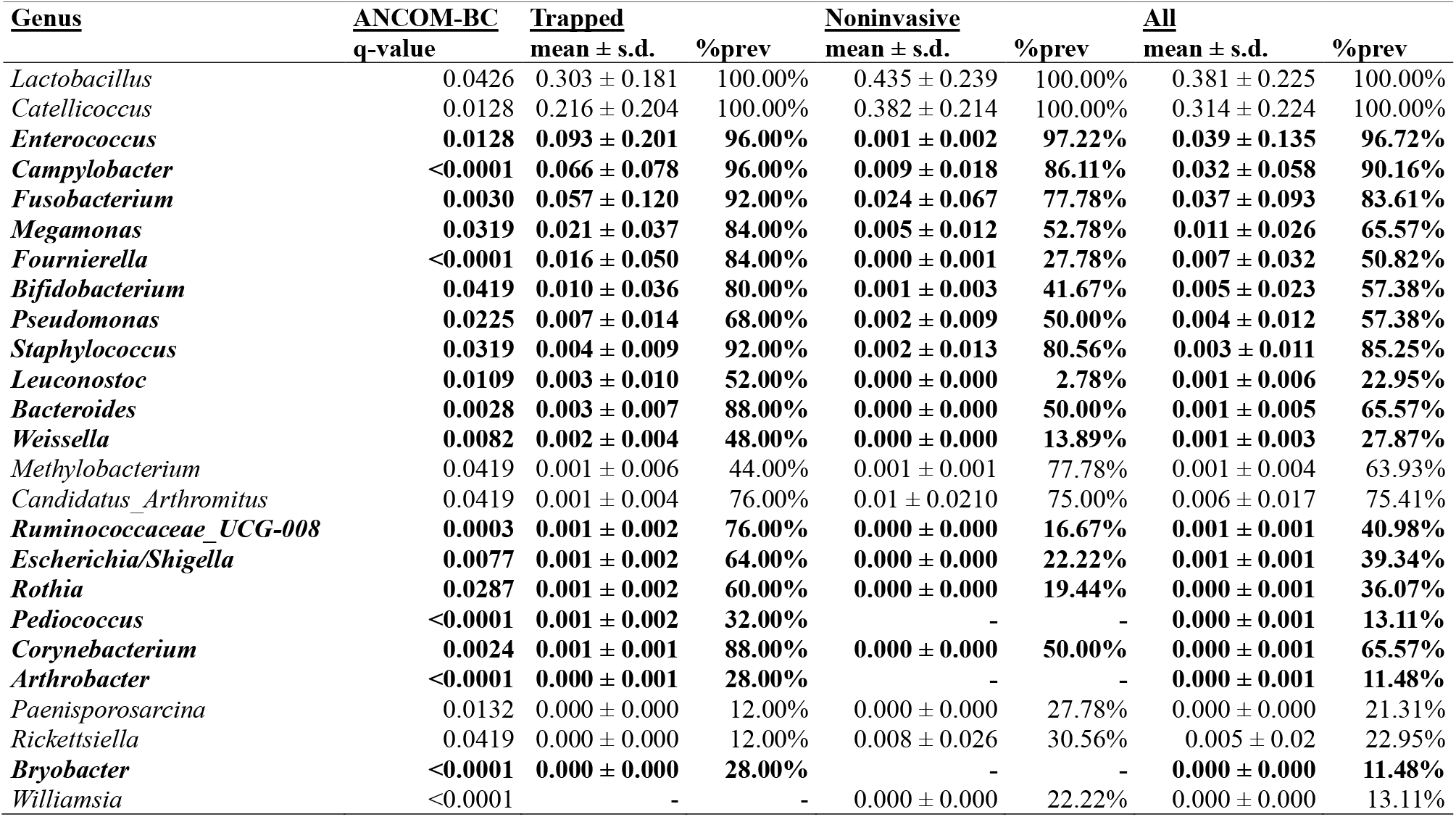
Significant genera from differential abundance analysis. The significant genera identified using ANCOM-BC (and presented in Fig. 4c) are presented here with their mean relative abundance across the groups as well as their per-group prevalence. Data for the entire set of samples (All) is also presented. Data are ordered by mean relative abundance of the trapped group. Bolded rows are enriched in the trapped samples and rows that are not bolded are enriched in the noninvasive samples.

## References

Banerjee, S., Schlaeppi, K., & van der Heijden, M. G. A. (2018). Keystone taxa as drivers of microbiome structure and functioning. Nature Reviews Microbiology 2018 16:9, 16(9), 567–576. https://doi.org/10.1038/s41579-018-0024-1

Battaglia, T. (2021). btools: A suite of R function for all types of microbial diversity analyses. R package version 0.0.1.

Beja-Pereira, A., Oliveira, R., Alves, P. C., Schwartz, M. K., & Luikart, G. (2009). Advancing ecological understandings through technological transformations in noninvasive genetics. Molecular Ecology Resources, 9(5), 1279–1301. https://doi.org/10.1111/J.1755-0998.2009.02699.X

Blomberg, E. J., Davis, S. B., Mangelinckx, J., & Sullivan, K. (2018). Detecting capture-related mortality in radio-marked birds following release. Avian Conservation and Ecology, 13(1). https://doi.org/10.5751/ACE-01147-130105

Cerutti, F., Modesto, P., Rizzo, F., Cravero, A., Jurman, I., Costa, S., Giammarino, M., Mandola, M. L., Goria, M., Radovic, S., Cattonaro, F., Acutis, P. L., & Peletto, S. (2018). The microbiota of hematophagous ectoparasites collected from migratory birds. PLoS ONE, 13(8), e0202270. https://doi.org/10.1371/journal.pone.0202270

Choi, O. N., Corl, A., Wolfenden, A., Lublin, A., Ishaq, S. L., Turjeman, S., Getz, W. M., Nathan, R., Bowie, R. C. K., & Kamath, P. L. (2021). High-throughput sequencing for examining *Salmonella* prevalence and pathogen—microbiota relationships in barn swallows. Frontiers in Ecology and Evolution, 9, 681. https://doi.org/10.3389/fevo.2021.683183

Čížková, D., Ďureje, Ľ., Piálek, J., & Kreisinger, J. (2021). Experimental validation of small mammal gut microbiota sampling from faeces and from the caecum after death. Heredity, 127(2), 141–150. https://doi.org/10.1038/s41437-021-00445-6

Collins, S. M., & Bercik, P. (2009). The relationship between intestinal microbiota and the central nervous system in normal gastrointestinal function and disease. Gastroenterology, 136(6), 2003–2014. https://doi.org/10.1053/J.GASTRO.2009.01.075

Comizzoli, P., & Power, M. (2019). Reproductive microbiomes in wild animal species: A new dimension in conservation biology. In Advances in Experimental Medicine and Biology (Vol. 1200, pp. 225–240). Adv Exp Med Biol. https://doi.org/10.1007/978-3-030-23633-5_8

Davis, N. M., Proctor, Di. M., Holmes, S. P., Relman, D. A., & Callahan, B. J. (2018). Simple statistical identification and removal of contaminant sequences in marker-gene and metagenomics data. Microbiome, 6(1), 1–14. https://doi.org/10.1186/s40168-018-0605-2

Dennis, T. E., & Shah, S. F. (2012). Assessing acute effects of trapping, handling, and tagging on the behavior of wildlife using GPS telemetry: A case study of the common brushtail possum. Journal of Applied Animal Welfare Science, 15(3), 189–207. https://doi.org/10.1080/10888705.2012.683755

Escallón, C., Belden, L. K., & Moore, I. T. (2019). The cloacal microbiome changes with the breeding season in a wild bird. Integrative Organismal Biology, 1(1), 1–16. https://doi.org/10.1093/IOB/OBY009

Gibson, K. M., Nguyen, B. N., Neumann, L. M., Miller, M., Buss, P., Daniels, S., Ahn, M. J., Crandall, K. A., & Pukazhenthi, B. (2019). Gut microbiome differences between wild and captive black rhinoceros - implications for rhino health. Scientific Reports, 9(1). https://doi.org/10.1038/S41598-019-43875-3

Guo, W., Ren, K., Ning, R., Li, C., Zhang, H., Li, D., Xu, L., Sun, F., & Dai, M. (2020). Fecal microbiota transplantation provides new insight into wildlife conservation. Global Ecology and Conservation, 24, e01234. https://doi.org/10.1016/J.GECCO.2020.E01234

Hartup, B. K., Schneider, L., Michael Engels, J., Hayes, M. A., & Barzen, J. A. (2014). Capture of sandhill cranes using alpha-chloralose: A 10-year follow-up. In Journal of Wildlife Diseases (Vol. 50, Issue 1, pp. 143–145). Allen Press. https://doi.org/10.7589/2013-06-140

Hayes, M. A., Hartup, B. K., Pittman, J. M., & Barzen, J. A. (2003). Capture of sandhill cranes using alpha-chloralose. Journal of Wildlife Diseases, 39(4), 859–868. https://doi.org/10.7589/0090-3558-39.4.859

Knutie, S. A., & Gotanda, K. M. (2018). A non-invasive method to collect fecal samples from wild birds for microbiome studies. Microbial Ecology, 76(4), 851–855. https://doi.org/10.1007/s00248-018-1182-4

Kolde, R. (2019). pheatmap: Pretty Heatmaps. R package version 1.0.12.

Koskey, A. M., Fisher, J. C., Traudt, M. F., Newton, R. J., & McLellan, S. L. (2014). Analysis of the gull fecal microbial community reveals the dominance of *Catellicoccus marimammalium* in relation to culturable enterococci. Applied and Environmental Microbiology, 80(2), 757–765. https://doi.org/10.1128/AEM.02414-13/ASSET/00328B17-6BD3-4DC2-9EB9-06F126C32A44/ASSETS/GRAPHIC/ZAM9991050630003.JPEG

Lahti, L., & Shetty, S. (2019). microbiome R package.

Larsson, J., & Gustafsson, P. (2018). A case study in fitting area-proportional euler diagrams with ellipses using eulerr. 84–91. https://doi.org/10.2/JQUERY.MIN.JS

Lin, H., & Peddada, S. das. (2020). Analysis of compositions of microbiomes with bias correction. Nature Communications 2020 11:1, 11(1), 1–11. https://doi.org/10.1038/s41467-020-17041-7

Lyte, J. M., Keane, J., Eckenberger, J., Anthony, N., Shrestha, S., Marasini, D., Daniels, K. M., Caputi, V., Donoghue, A. M., & Lyte, M. (2021). Japanese quail (*Coturnix japonica*) as a novel model to study the relationship between the avian microbiome and microbial endocrinology-based host-microbe interactions. Microbiome, 9(1), 1–24. https://doi.org/10.1186/s40168-020-00962-2

McMurdie, P. J., & Holmes, S. P. (2013). phyloseq: An R package for reproducible interactive analysis and graphics of microbiome census data. PLoS ONE, 8(4), e61217. https://doi.org/10.1371/journal.pone.0061217

Niederwerder, M. C. (2018). Fecal microbiota transplantation as a tool to treat and reduce susceptibility to disease in animals. Veterinary Immunology and Immunopathology, 206, 65. https://doi.org/10.1016/J.VETIMM.2018.11.002

Noguera, J. C., Aira, M., Pérez-Losada, M., Domínguez, J., & Velando, A. (2018). Glucocorticoids modulate gastrointestinal microbiome in a wild bird. Royal Society Open Science, 5(4). https://doi.org/10.1098/RSOS.171743

Oksanen, J., Blanchet, F. G., Kindt, R., Legendre, P., Minchin, P. R., O’Hara, R. B., Simpson, G. L., Solymos, P., Stevens, M. H. H., & Wagner, H. H. (2018). vegan: Commnity ecology package. R package version 2.5-3 (2.5.3). https://cran.r-project.org/package=vegan

Oliveira, B. C. M., Murray, M., Tseng, F., & Widmer, G. (2020). The fecal microbiota of wild and captive raptors. Animal Microbiome, 2(1), 1–9. https://doi.org/10.1186/s42523-020-00035-7

Pannoni, S. B., Proffitt, K. M., & Holben, W. E. (2022). Non-invasive monitoring of multiple wildlife health factors by fecal microbiome analysis. Ecology and Evolution, 12(2), e8564. https://doi.org/10.1002/ECE3.8564

Peixoto, R. S., Harkins, D. M., & Nelson, K. E. (2021). Advances in Microbiome Research for Animal Health. Annual Review of Animal Biosciences, 9, 289–311. https://doi.org/10.1146/ANNUREV-ANIMAL-091020-075907

Pekarsky, S., Corl, A., Turjeman, S., Kamath, P. L., Getz, W. M., Bowie, R. C. K., Markin, Y., & Nathan, R. (2021). Drivers of change and stability in the gut microbiota of an omnivorous avian migrant exposed to artificial food supplementation. Molecular Ecology, 30(19), 4723–4739. https://doi.org/10.1111/MEC.16079

Risely, A. (2020). Applying the core microbiome to understand host–microbe systems. In Journal of Animal Ecology (Vol. 89, Issue 7, pp. 1549–1558). John Wiley & Sons, Ltd. https://doi.org/10.1111/1365-2656.13229

Russello, M. A., Waterhouse, M. D., Etter, P. D., & Johnson, E. A. (2015). From promise to practice: Pairing non-invasive sampling with genomics in conservation. PeerJ, 2015(7), e1106. https://doi.org/10.7717/PEERJ.1106/SUPP-1

Serbanescu, M. A., Mathena, R. P., Xu, J., Santiago-Rodriguez, T., Hartsell, T. L., Cano, R. J., & Mintz, C. D. (2019). General anesthesia alters the diversity and composition of the intestinal microbiota in mice. Anesthesia and Analgesia, 129(4), e126–e129. https://doi.org/10.1213/ANE.0000000000003938

Snel, J., Heinen, P. P., Blok, H. J., Carman, R. J., Duncan, A. J., Allen, P. C., & Collins, M. D. (1995). Comparison of 16S rRNA sequences of segmented filamentous bacteria isolated from mice, rats, and chickens and proposal of “*Candidatus Arthromitus*.” International Journal of Systematic Bacteriology, 45(4), 780–782. https://doi.org/10.1099/00207713-45-4-780

Spotswood, E. N., Goodman, K. R., Carlisle, J., Cormier, R. L., Humple, D. L., Rousseau, J., Guers, S. L., & Barton, G. G. (2012). How safe is mist netting? Evaluating the risk of injury and mortality to birds. Methods in Ecology and Evolution, 3(1), 29–38. https://doi.org/10.1111/j.2041-210X.2011.00123.x

Stothart, M. R., Bobbie, C. B., Schulte-Hostedde, A. I., Boonstra, R., Palme, R., Mykytczuk, N. C. S., & Newman, A. E. M. (2016). Stress and the microbiome: Linking glucocorticoids to bacterial community dynamics in wild red squirrels. Biology Letters, 12(1). https://doi.org/10.1098/rsbl.2015.0875

Sutherland, W. J., Newton, I., & Green, R. E. (2004). Bird Ecology and Conservation: A Handbook of Techniques. Oxford University Press.

Trevelline, B. K., Fontaine, S. S., Hartup, B. K., & Kohl, K. D. (2019). Conservation biology needs a microbial renaissance: a call for the consideration of host-associated microbiota in wildlife management practices. Proceedings of the Royal Society B, 286(1895). https://doi.org/10.1098/RSPB.2018.2448

Turjeman, S., & Koren, O. (2021). Using the microbiome in clinical practice. Microbial Biotechnology. https://doi.org/10.1111/1751-7915.13971

Vandeputte, D., Falony, G., Vieira-Silva, S., Tito, R. Y., Joossens, M., & Raes, J. (2016). Stool consistency is strongly associated with gut microbiota richness and composition, enterotypes and bacterial growth rates. Gut, 65(1), 57–62. https://doi.org/10.1136/GUTJNL-2015-309618

